# Inhibition of Inflammatory Osteoclasts Enhances CGRP^+^TrkA^+^ Signaling and Accelerates Callus Remodeling in Osteoporotic Fractures

**DOI:** 10.1101/2024.05.21.595143

**Authors:** Yuexia Shu, Zhenyu Tan, Zhen Pan, Yujie Chen, Jielin Wang, Jieming He, Jia Wang, Yuan Wang

## Abstract

Impaired callus remodeling significantly contributes to the delayed healing of osteoporotic fractures; however, the underlying mechanisms remain unclear. Sensory neuronal signaling plays a crucial role in bone repair. In this study, we demonstrate that in ovariectomized (OVX) mice, the loss of CGRP^+^TrkA^+^ sensory neuronal signaling during callus remodeling correlates with increased Cx3cr1^+^iOCs expression within the bone callus. Conditional knockout of Cx3cr1^+^iOCs restored CGRP^+^TrkA^+^ sensory neuronal, enabling normal callus remodeling progression. Mechanistically, we further demonstrate that Cx3cr1^+^iOCs secrete seme3A in the osteoporotic fracture repair microenvironment, inhibiting CGRP^+^TrkA^+^ sensory neurons’ axonal regeneration and suppressing nerve-bone signaling exchange, thus hindering bone remodeling. Lastly, in human samples, we observed an association between the loss of CGRP^+^TrkA^+^ sensory neuronal signaling and increased expression of Cx3cr1^+^iOCs. In conclusion, enhancing CGRP^+^TrkA^+^ sensory nerve signaling by inhibiting Cx3cr1^+^iOCs activity presents a potential strategy for treating delayed healing in osteoporotic fractures.

## Introduction

With the aging of the population, the incidence of osteoporotic fractures increases, and delayed bone healing is complicated(Compston et al., 2019). Delayed healing of osteoporotic fractures not only results in long-term bed rest for elderly patients, but also increases both mortality risks and healthcare costs(Clynes et al., 2020; Lewiecki et al., 2019). Current treatments like Teriparatide, which promotes bone formation, and bisphosphonates, that inhibit bone resorption, reduce fracture risk but do not accelerate fracture healing(Qaseem et al., 2023). Due to the lack of understanding of the mechanisms behind this healing process, an effective treatment strategy has yet to be found.

Sensory nerve fibers transmit nociceptive signals, particularly evident following bone fractures(Mantyh et al., 2011; Neychev et al., 2020). Numerous studies indicate that calcitonin gene-related peptide (CGRP^+^) sensory nerves have an evolutionarily conserved, crucial role in organ formation and tissue regeneration(Tomlinson et al., 2016). Recent research has shown that the TrkA signaling of CGRP^+^ sensory nerves in bones is essential for fracture repair(Li et al., 2019). Nerve growth factor initiates fracture repair by activating the TrkA signaling pathway. However, it remains uncertain whether TrkA signaling in CGRP+ sensory nerves also influence fractures repair in osteoporotic fractures.

Fracture healing typically includes three phases: inflammatory, callus formation, and callus remodeling(Bahney et al., 2019). Callus remodeling, the final repair stage, is critical for bone to regain normal function. Osteoclasts play a critical regulatory role during this process(Boyle et al., 2003). Normally, osteoclasts work with other cells through a "phagocytic resorption" mechanism to rebuild bone and transform dense calluses into organized trabecular structures. However, in pathological states, changes in osteoclast differentiation or resorption activity are closely linked to diseases like osteoporosis(Dou et al., 2022). Studies using the ovariectomized mouse (OVX) model show that callus remodeling is impaired, and trabecular formation is delayed(Chen et al., 2021). Recent research has found that in osteoporosis, some osteoclasts, termed inflammatory osteoclasts (iOCs), show high expression of Cx3cr1(Madel et al., 2020). Functional abnormalities in these Cx3cr1^+^iOCs may disrupt normal bone remodeling and delay fracture healing in pathological states. However, further investigation is required to elucidate the specific roles and mechanisms of Cx3cr1^+^iOCs in delayed healing associated with osteoporosis.

This study explored the pathological mechanisms obstructing bone remodeling in osteoporotic fractures. During the critical callus remodeling phase, we observed that elevated expression of Cx3cr1^+^iOCs inhibits CGRP^+^TrkA^+^ sensory neural signaling, thereby contributing significantly to delayed callus remodeling. Eliminating either Sema3A secreted by Cx3cr1^+^iOCs or the Cx3cr1^+^iOCs themselves reversed this inhibition, restoring normal callus remodeling. Furthermore, analysis of human samples revealed a correlation between diminished CGRP^+^TrkA^+^ sensory neural signaling and increased expression of Cx3cr1^+^iOCs. These findings enhance our understanding of the pathological mechanisms underlying delayed healing in osteoporotic fractures.

## Results

### Upregulation of Cx3cr1^+^iOCs in the bone callus is associated with diminished trabecular formation during the remodeling of osteoporotic fractures

Osteoporotic fractures frequently occur in postmenopausal women. To mimic this clinical scenario, bilateral ovariectomy was performed on 10-week-old WT mice, followed by the induction of femoral fractures 8 weeks after OVX surgery (Fig. 1 A). Micro-computed tomography (Micro-CT) was utilized to assess bone mass in OVX mice. The establishment of an osteoporosis mouse model was successfully confirmed by the results (Fig. S1, A-D). To explore the effects of osteoporosis on fracture healing, Micro-CT analyses were conducted on OVX and control mice at various post-fracture time points. At the same post-fracture time points, Micro-CT analysis revealed that the OVX group showed significantly less callus formation and remodeling than the control group (Fig. 1 A and Fig. S1, E and F). Micro-CT quantitative analyses indicated that, in OVX mice, the bone volume to total volume (BV/TV) ratios decreased by 49% and 47% at 2- and 4-weeks post-fracture, respectively (Fig. 1 F). Additionally, OVX mice demonstrated 44% and 34% reduction in the number (Tb.N) and thickness (Tb.Th) of trabeculae in the callus, respectively, compared to the control group during the same periods (Fig. 1, G and H). Hematoxylin-eosin (HE) staining revealed significant cellular accumulation at the fracture sites in control mice, forming a cartilaginous callus early in the healing process (Fig. 1 C). This callus then successfully mineralized and remodeled into trabeculae (Fig. 1 C). In contrast, the OVX group showed inhibited early callus formation, leading to delayed trabecular mineralization in later stages (Fig. 1 C). Saffron O-robust green (SO-FG) and Masson staining demonstrated delayed callus formation and mineralization in OVX mice (Fig. 1 D, Fig. S1 G).

**Figure. 1.**
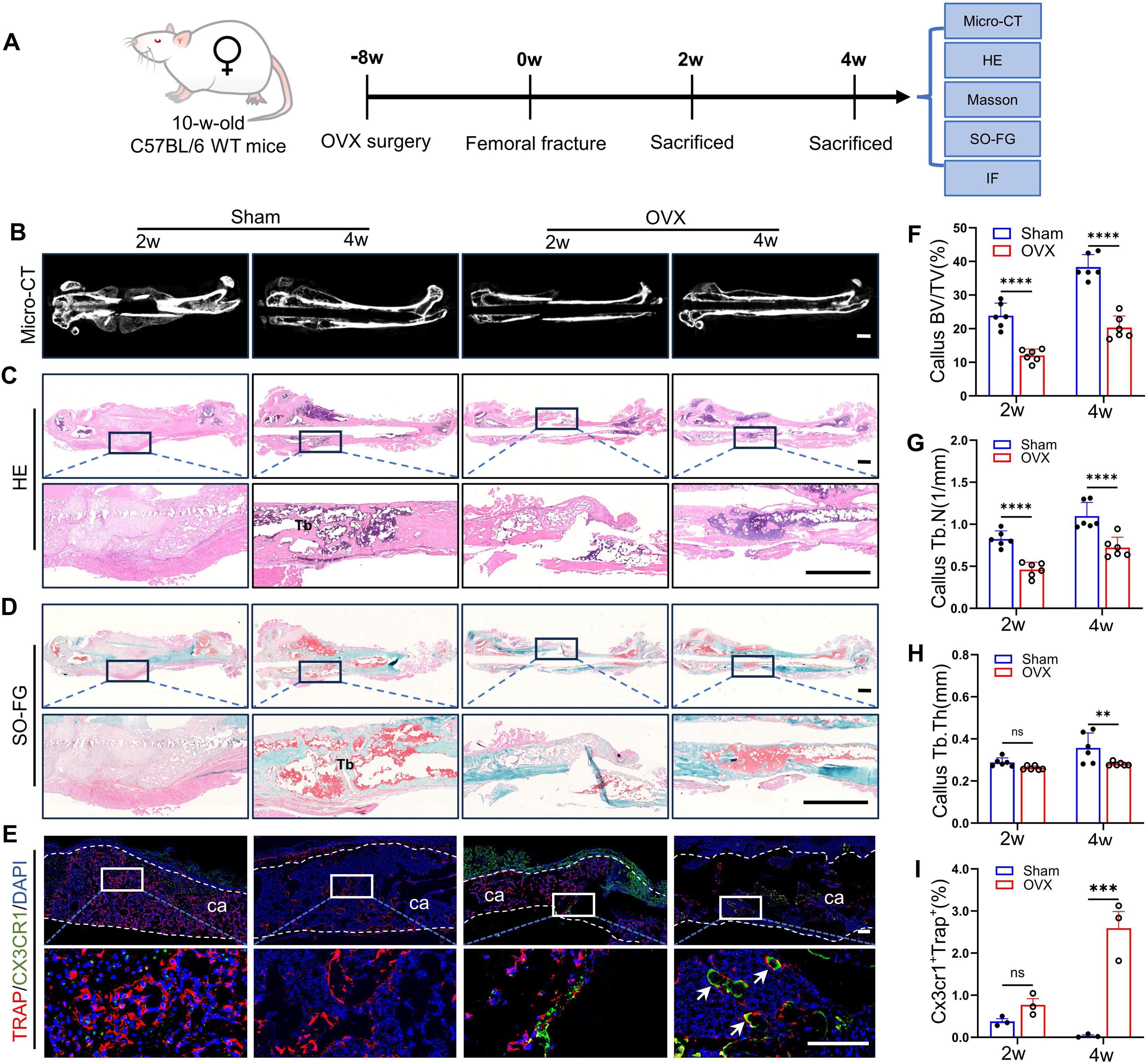
Upregulation of Cx3cr1^+^iOCs in the bone callus is associated with diminished trabecular formation during the remodeling of osteoporotic fractures. (A) Schematic of the experimental design to observe the role of Cx3cr1^+^iOCs in callus remodeling during osteoporotic fractures. (B) Representative micro-CT images of femoral fractures at 2- and 4-weeks post-fracture. Scale bar: 1 mm. (C-D) Representative HE (C) and SO-FG) (D) stained images of femoral fractures in Sham and OVX mice at 2- and 4-weeks post-fracture. Upper panels show global views; lower panels provide close-up views of the fracture sites. Tb: trabecular bone. Scale bar: 1 mm. (E) Representative Cx3cr1 and Trap coimmunostaining images of calluses from Sham and OVX mice at 2- and 4-weeks post-fracture. ca: callus. White dashed lines outline the callus boundaries, and white arrows point to Cx3cr1^+^Trap^+^ cells (Cx3cr1^+^iOCs). Scale bar: 200 μm. (F-H) Micro-CT quantitative analysis of the callus from (B). Parameters include bone volume fraction (BV/TV), trabecular thickness (Tb.Th), and trabecular number (Tb.N). n = 6. (I) Quantification of Cx3cr1^+^Trap^+^-positive osteoclasts (Cx3cr1^+^iOCs) in the callus from (E). n = 3. Data presented as mean ± SEM. *P < 0.05, **P < 0.01, ***P < 0.001, ****P < 0.0001.

Osteoclast activity is essential for callus remodeling, as it regulates callus remodeling through bone resorption(Teitelbaum, 2007). Tartrate-resistant acid phosphatase is a marker of osteoclast function(Minkin, 1982). Double immunofluorescence staining for Cx3cr1^+^Trap^+^ revealed that while WT mice predominantly exhibited Trap^+^ positive cells within the calluses (Fig.1 E), the number of Cx3cr1^+^Trap^+^ cells significantly increased in OVX mice during critical phases of callus formation and trabecular development (Fig.1, E and I). Thus, in the context of osteoporotic fracture repair, the suppression of callus remodeling and trabecular formation correlates with an elevated expression of Cx3cr1^+^iOCs at the fracture sites.

### Conditional knockout of Cx3cr1^+^iOCs accelerates callus remodeling in OVX mice

To further validate the role of Cx3cr1^+^iOCs in callus remodeling during osteoporotic fractures, we developed a specific conditional Cx3cr1^+^iOCs knockout mouse model (Fig.2 A). Micro-CT analysis revealed that callus remodeling and trabecular formation were significantly enhanced in Cx3cr1-Cre; DTR^fl/fl^ OVX mice compared to WT OVX mice (Fig. 2B and Fig.S2 A). Quantitative Micro-CT analysis showed a 28.6% increase in callus BV/TV in Cx3cr1^+^iOCs-deficient OVX mice compared to WT OVX mice, with no significant changes in the Sham group (Fig.2 F). Additionally, in Cx3cr1^+^iOCs-deficient OVX mice, the Tb.N in the callus increased by 55%, and the Tb.Th decreased by 16% (Fig. 2, G and H). These results suggest that the absence of CX3CR1-iOCs promotes increased trabecular formation and accelerates callus remodeling.

**Figure. 2.**
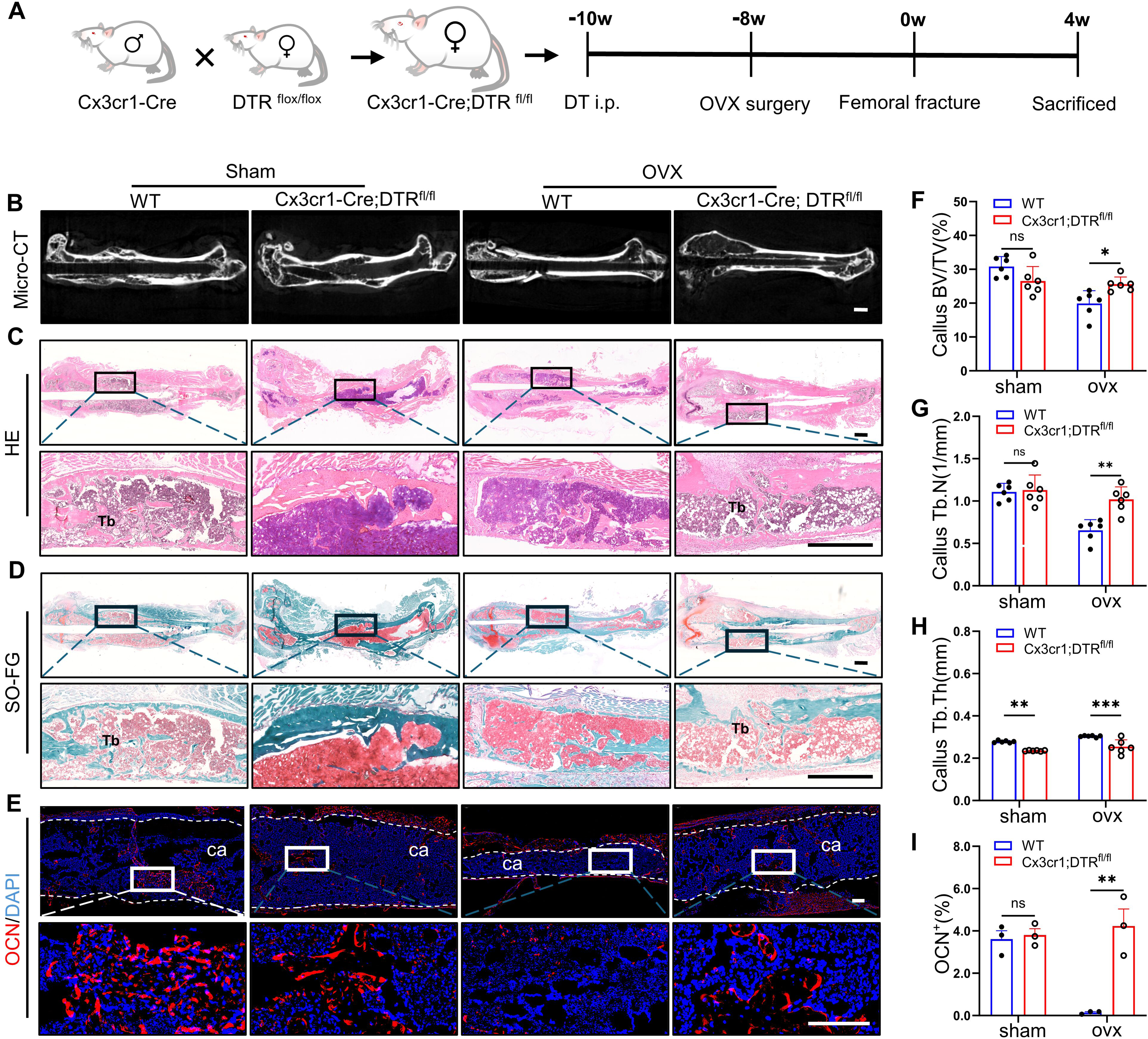
Conditional knockout of Cx3cr1^+^iOCs accelerates callus remodeling in OVX mice. (A) Schematic of the experimental design to assess the effects of Cx3cr1^+^iOCs deficiency on callus remodeling in osteoporotic fractures. (B) Representative Micro-CT images of femoral fractures in Cx3cr1-Cre; DTR^fl/fl^ mice and WT littermates at 4 weeks post-fracture. Scale bar: 1 mm. (C-D) Representative HE (C) and SO-FG) (D) stained images of femoral fractures in Cx3cr1-Cre; DTR^fl/fl^ mice and WT littermates at 4 weeks post-fracture. Upper panels display global views; lower panels provide close-ups of the fracture sites. Tb: trabecular bone. Scale bar: 1 mm. (E) Immunofluorescence staining images showing osteocalcin (OCN) in calluses of Cx3cr1-Cre; DTR^fl/fl^ mice and WT littermates at 4 weeks post-fracture. ca: callus. White dashed lines outline the callus boundaries. Scale bar: 200 μm. (F-H) Micro-CT quantitative analysis of the callus from (B). Parameters include bone volume fraction (BV/TV), trabecular thickness (Tb.Th), and trabecular number (Tb.N). n = 6. (I) Quantitative analysis of OCN fluorescence intensity in callus from (E). n = 3. Data presented as mean ± SEM. *P < 0.05, **P < 0.01, ***P < 0.001 and ****P < 0.0001.

Consistent with Micro-CT findings, HE staining showed an increased number of trabeculae in the bone callus of Cx3cr1-Cre; DTR^fl/fl^ OVX mice compared to their littermate controls (Fig.2 C). SO-FG and Masson staining both revealed a higher number of newly formed trabeculae in the bone callus of Cx3cr1-Cre; DTR^fl/fl^ OVX mice (Fig.2 A and Fig.S2 B). Osteocalcin (OCN) plays a pivotal role in the mineralization of the bone callus (Komori, 2020). Immunofluorescence staining for OCN showed elevated expression levels in the bone callus of Cx3cr1-Cre; DTR^fl/fl^ OVX mice (Fig.2, E and I). However, no such differences were observed in the Sham group (Fig.2, E and I). In summary, the conditional knockout of Cx3cr1^+^iOCs may counteract the trend of impaired bone callus remodeling observed in osteoporotic fractures.

### In osteoporotic fractures, attenuation of CGRP^+^TrkA^+^ signaling correlates with upregulated Cx3cr1^+^iOCs expression in the callus, and the conditional knockout of Cx3cr1^+^iOCs enhances CGRP^+^TrkA^+^ signaling

Under physiological conditions, the growth of CGRP^+^TrkA^+^ sensory nerves accompanies the entire fracture repair process(Li et al., 2019). We hypothesize that increased expression of Cx3cr1^+^iOCs in osteoporotic fractures may inhibit neural signal transmission within the callus. To test this hypothesis, immunofluorescence techniques were used to examine sensory nerve sprouting within the callus. Our findings indicate minimal expression of sensory nerve signaling in the early stages of fracture repair, localized near the inner and outer membranes of the callus (Fig. S2 C). Consequently, we focused on observing the sensory nerve signaling during the bone callus remodeling. During callus remodeling, the expression of TUBB3 protein in the callus of Cx3cr1-Cre; DTR^fl/fl^ OVX mice was found to be 2.5-fold higher compared to WT OVX mice, while no significant difference was observed in sham mice (Fig. 3, A and B). Given the predominance of CGRP+ nerves among sensory nerve fibers in the callus, our investigation shifted to exploring the role of CGRP signaling (Li et al., 2024). Following this, the expression of CGRP in the callus was evaluated. Consistent with previous findings, CGRP protein levels were 77-fold higher in the callus of Cx3cr1-Cre; DTR^fl/fl^ OVX mice compared to WT OVX mice (Fig. 3, C and D). Additionally, TrkA immunofluorescence was used to examine its expression in the callus. Surprisingly, TrkA protein levels in the callus of Cx3cr1-Cre; DTR^fl/fl^ OVX mice were nearly 124-fold higher than in WT OVX mice (Fig. 3, E and F). In summary, reduced CGRP^+^TrkA^+^ signaling at the fracture site correlates with impaired callus remodeling in osteoporotic fractures, while the knockdown of Cx3cr1^+^iOCs enhances CGRP^+^TrkA^+^ signaling by sensory nerves.

**Figure. 3.**
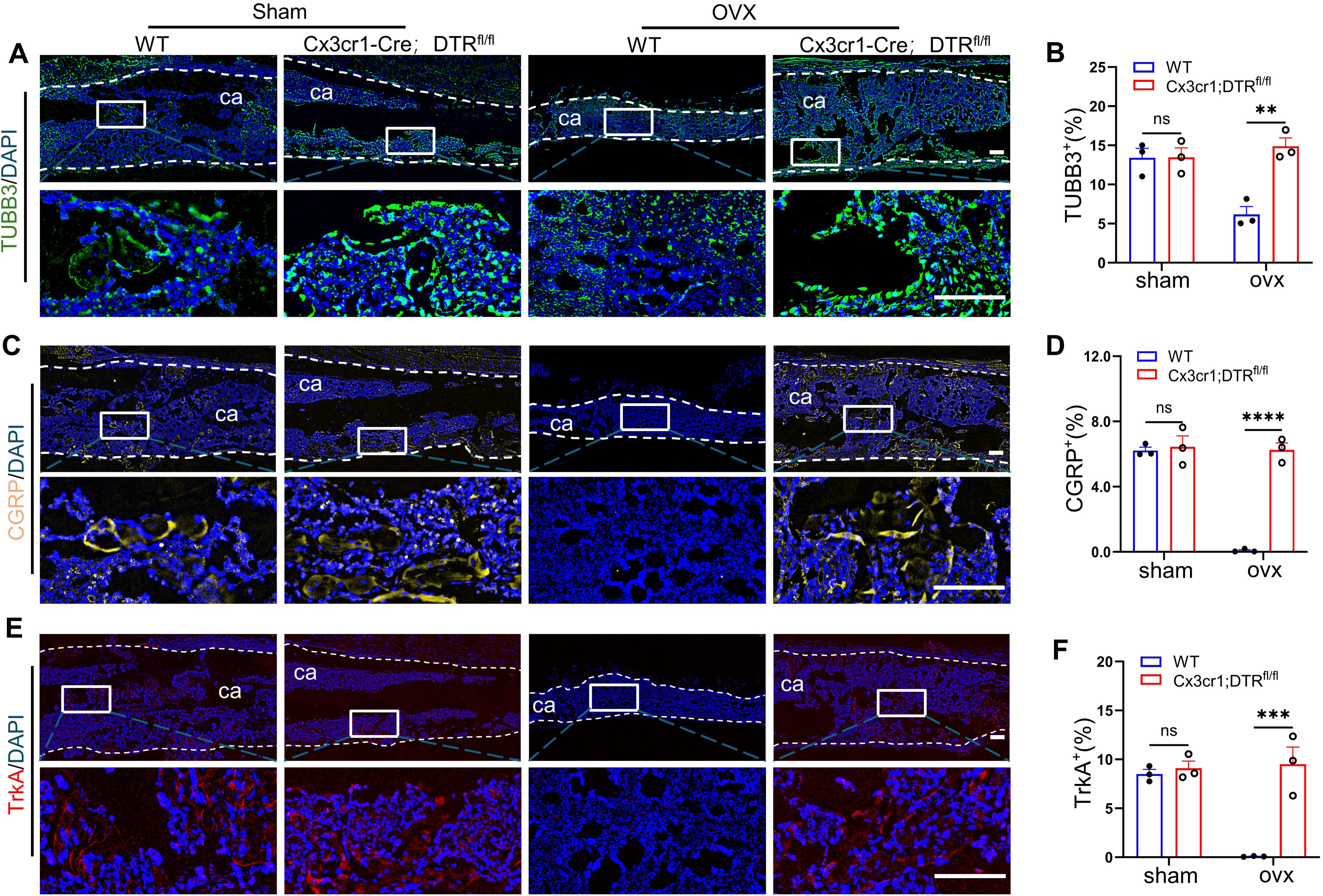
In osteoporotic fractures, attenuation of CGRP^+^TrkA^+^ signaling correlates with upregulated Cx3cr1^+^iOCs expression in the callus, and the conditional knockout of Cx3cr1^+^iOCs enhances CGRP^+^TrkA^+^ signaling. (A-B) Immunofluorescence staining images and quantification of TUBB3 protein levels in calluses from Cx3cr1-Cre; DTR^fl/fl^ mice and WT littermates at 4 weeks post-fracture. (C-D) Immunofluorescence staining images and quantification of CGRP protein levels in calluses from Cx3cr1-Cre; DTR^fl/fl^ mice and WT littermates at 4 weeks post-fracture. (E-F) Immunofluorescence staining images and quantification of TrkA protein levels in calluses from Cx3cr1-Cre; DTR^fl/fl^ mice and WT littermates at 4 weeks post-fracture. White dashed lines outline the callus boundaries. ca: callus. Scale bar: 200 μm. n = 3 per group; data presented as mean ± SEM. *P < 0.05, **P < 0.01, ***P < 0.001 and ****P < 0.0001.

### Sema3A secreted by cx3cr1^+^iOCs inhibits CGRP^+^TrkA^+^ sensory nerve signaling via GSK3β-Akt pathway in vitro

We further investigated how Cx3cr1^+^iOCs influence callus remodeling via the CGRP^+^TrkA^+^ signaling pathway. Considering that osteoclasts derive from the monocyte-macrophage lineage (Boyle et al., 2003), we analyzed the proportion of Cx3cr1^+^Ly6C^+^ cells in mouse bone marrow by using flow cytometry (Fig. 4, A and B and Fig. S3 A). The results demonstrated a significant increase in Cx3cr1^+^Ly6C^+^ cells in OVX mice compared to WT mice (Fig. 4, A and B and Fig. S3 A). We hypothesize that Cx3cr1^+^iOCs secrete inhibitory signals that impede the regeneration of CGRP^+^TrkA^+^ sensory nerve fibers in the callus. Semaphorins, widely expressed in bone, inhibit axonal nerve growth (Kang and Kumanogoh, 2013). We then assessed semaphorin expression in the callus using immunofluorescence staining, which revealed a notable increase in the expression of Sema3A secreted by Cx3cr1^+^iOCs (Fig. 4, C and D and Fig. S3 B). To elucidate the impact of Cx3cr1^+^iOCs-derived Sema3A on sensory nerve signaling, we engineered Sema3A shRNA to silence Sema3A in these cells and co-cultured them with dorsal root ganglia (DRG) (Fig. 4 E). TUBB3 immunofluorescence staining was performed to access axonal regeneration of DRG. TUBB3 protein expression was significantly reduced in the OVX group compared to the sham group and increased significantly after silencing Sema3A in Cx3cr1^+^iOCs (Fig. 4, F and G). Western blot analysis confirmed that silencing Sema3A substantially increased CGRP and the phosphorylation of TrkA in DRG cells (Fig. 4, H-J). Conversely, Sema3A overexpression in the OVX group markedly reduced CGRP and p-TrkA expression in DRG cells (Fig. 4, H-J).

**Figure. 4.**
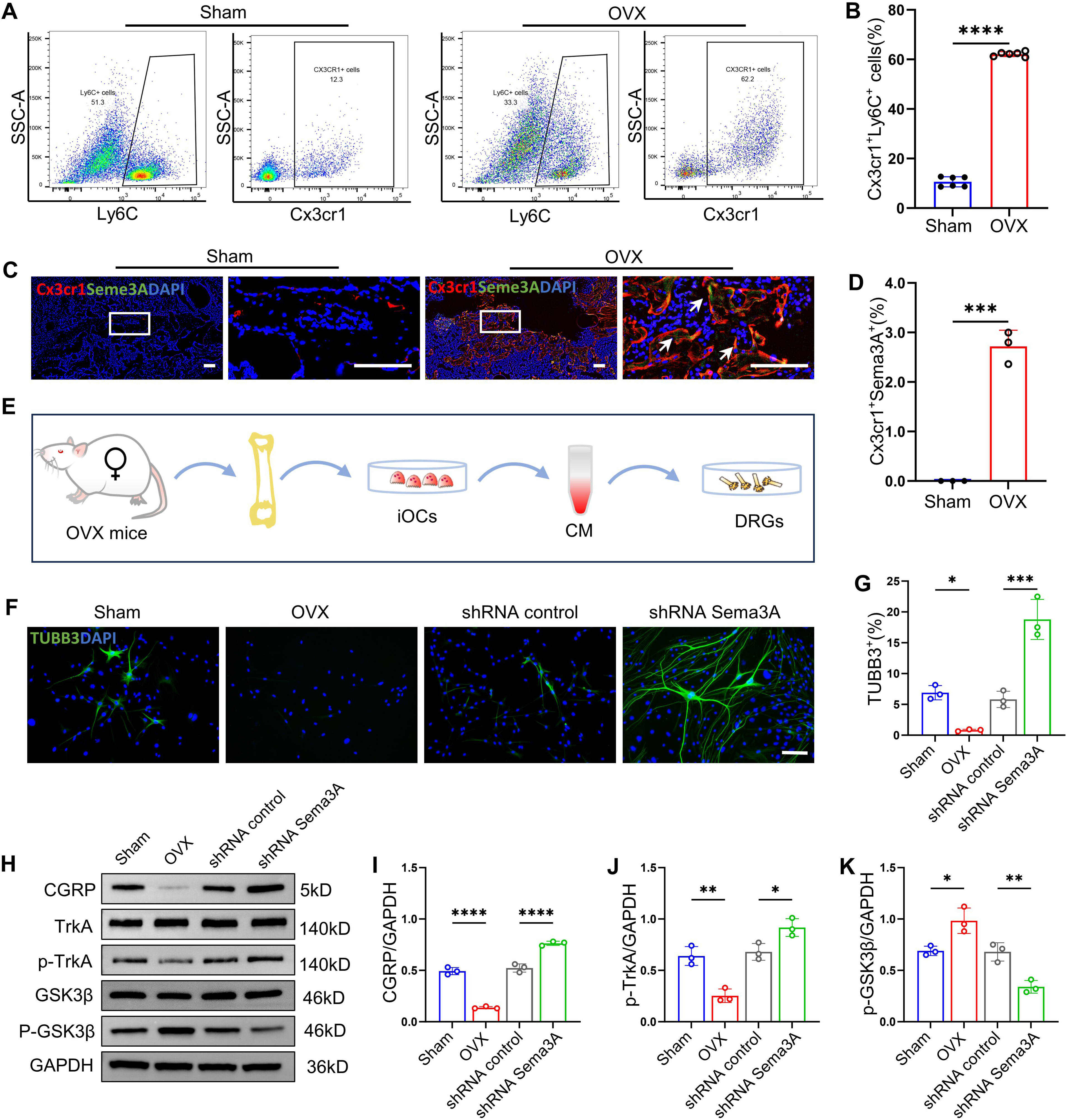
Sema3A secreted by Cx3cr1^+^iOCs inhibits CGRP^+^TrkA^+^ sensory nerve signaling via GSK3β-Akt pathway in vitro. (A-B) Flow cytometry analysis of Cx3cr1^+^Ly6C^+^ cells in bone marrow from sham and OVX mice. n=3. (C-D) Representative coimmunostaining images of Cx3cr1 and Sema3A and quantification in calluses from Sham and OVX mice at 4 weeks post-fracture.ca: callus. white arrows point to Cx3cr1^+^ Sema3A^+^ cells. Scale bar: 200 μm. n=3. (E) Schematic of DRGs co-cultured under under Cx3cr1^+^iOCs conditions. (F-G) Immunofluorescence staining and quantification of TUBB3 in Sham, OVX, shRNA control, and shRNA Sema3A groups. Scale bar: 50 μm. n=3. (H-K) Representative western blot images and quantification of CGRP, TrkA, p-TrkA, GSK3β, and p-GSK3β in DRGs treated with or without Sema3A from by cx3cr1^+^iOCs. n=3. Data presented as mean ± SEM. *P < 0.05, **P < 0.01, ***P < 0.001 and ****P < 0.0001.

Furthermore, to better understand how Sema3A from Cx3cr1^+^iOCs inhibits CGRP^+^TrkA^+^ signaling, we tested the expression and phosphorylation levels of GSK3β and Akt, including p-GSK3β and p-Akt. The results showed that the expression of p-GSK3βin OVX mice was increased compared with sham group, and the phosphorylation of GSK3βwas significantly reduced after silencing sema3A in Cx3cr1^+^iOCs. (Fig. 4, H and K and Fig. S3, C and E). The phosphorylation of AKT shows the opposite trend to p-GSK3β (Fig. S3, C and E). Consequently, we used LY2090314, a selective inhibitor of GSK3β. LY2090314 treatment blocked the inhibitory effect of Sema3A secreted by Cx3cr1^+^iOCs on CGRP^+^TrkA^+^ signaling. (Fig. S3, D, F and G). In summary, Sema3A derived from Cx3cr1^+^iOCs significantly inhibits CGRP^+^TrkA^+^ sensory nerve signaling via GSK3β-Akt pathway in vitro.

### Following the conditional knockout of Sema3A in Cx3cr1^+^iOCs, callus remodeling in OVX mice accelerated, accompanied by enhanced CGRP^+^TrkA^+^ signaling

To further investigate the role of Sema3A in Cx3cr1+iOCs during callus remodeling in osteoporotic fracture calluses, we developed Cx3cr1-Cre; Sema3A^fl/fl^ mice, featuring a targeted knockout of Sema3A in CX3CR1^+^iOCs (Fig. 5 A). HE staining demonstrated that both callus mineralization and trabecular formation were significantly enhanced in Cx3cr1-Cre; Sema3A^fl/fl^ OVX mice (Fig. 5 B). In contrast, extensive fibrous connections within the calluses were observed in the control group mice, with no significant mineralized trabeculae (Fig. 5 B). Additionally, SO-FG and Masson staining further confirmed significant acceleration in callus mineralization and trabecular formation in Cx3cr1-Cre; Sema3A^fl/fl^ OVX mice compared to their littermate controls (Fig. 5 C and Fig. S4 A). However, this change was not present in sham group.

**Figure. 5.**
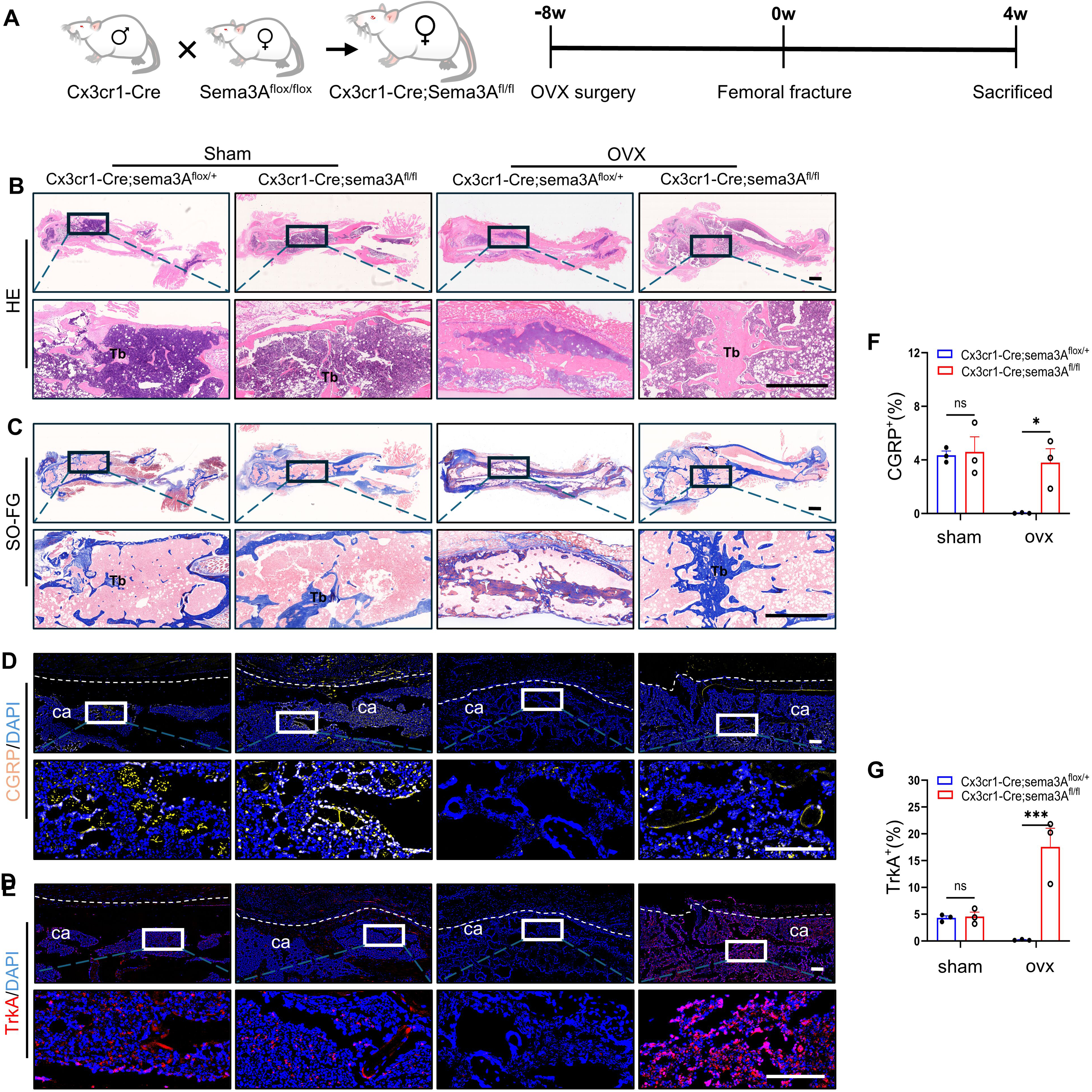
Following the conditional knockout of Sema3A in Cx3cr1^+^iOCs, callus remodeling in OVX mice accelerated, accompanied by enhanced CGRP^+^TrkA^+^ signaling. (A) Schematic of the experimental design to assess the effects of conditional deletion of Sema3A in Cx3cr1^+^iOCs on callus remodeling during osteoporotic fractures. (B-C) Representative HE (C) and Masson (D) stained images of femoral fractures in Cx3cr1-Cre; Sema3A^fl/fl^ OVX mice and their littermate controls at 4 weeks post-fracture. Upper panels display global views; lower panels show close-up views of the fracture sites. Tb: trabecular bone. Scale bar: 1 mm. (D and F) Immunofluorescence staining images and quantification of CGRP protein levels in calluses from Cx3cr1-Cre; Sema3A^fl/fl^ OVX mice and their littermate controls at 4 weeks post-fracture. ca: callus. White dashed lines outline the callus boundaries. Scale bar: 200 μm. n = 3. (E and G) Immunofluorescence staining images and quantification of TrkA protein levels in calluses from Cx3cr1-Cre; Sema3A^fl/fl^ OVX mice and their littermate controls at 4 weeks post-fracture. "Ca" denotes callus. White dashed lines outline the callus boundaries. Scale bar: 200 μm. n = 3. Data presented as mean ± SEM. *P < 0.05, **P < 0.01, ***P < 0.001, ****P < 0.0001.

Furthermore, we further verified the impact of Sema3A in Cx3cr1^+^iOCs on CGRP^+^TrkA^+^ signaling within the callus. CGRP immunofluorescence staining results indicated that the expression level of CGRP protein in the callus of Cx3cr1-Cre; Sema3A^fl/fl^ OVX mice increased by 108-fold (fig.5, D and F). Interestingly, abundant TrkA signaling was observed in the callus of Cx3cr1-Cre; Sema3A^fl/fl^ OVX mice (fig.5, D and F). The expression level of TrkA protein increased 90-fold compared to their littermate controls (fig.5, E and G). Similarly, this trend of change was not found in sham mice (fig.5, E and G). Additionally, the expression level of TUBB3 protein in the callus also showed a similar trend, showing a 26-fold rise (Fig. S4, B and C). In summary, the specific knockout of Sema3A in Cx3cr1^+^iOCs can reverse the loss of CGRP^+^TrkA^+^ signaling in osteoporotic fractures, suggesting that Cx3cr1^+^iOCs inhibit CGRP^+^TrkA^+^ signaling in the callus through the secretion of Sema3A, leading to impaired callus remodeling.

### Treatment with the Sema3A inhibitor TCS1105 reversed the delayed fracture healing observed in WT mice following OVX

Considering that the conditional knockout of Sema3A in Cx3cr1^+^iOCs promotes bone callus remodeling, we investigated Sema3A as a potential therapeutic target for osteoporotic fractures (Fig. 6 A). TCS1105, a Sema3A inhibitor, blocks the axonal growth defects in sensory neurons induced by Sema3A (van der Kooij et al., 2018).

**Figure. 6.**
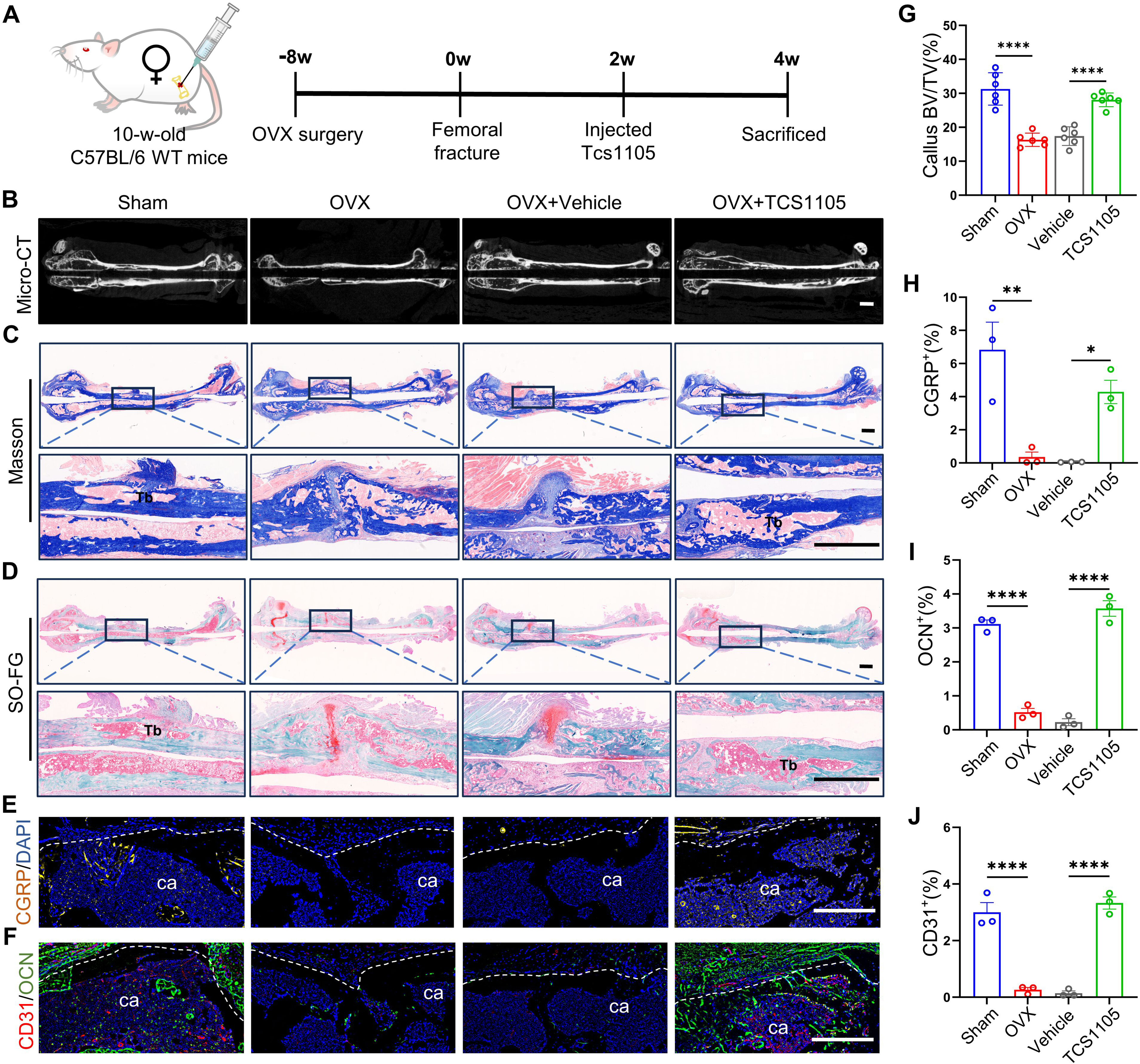
Treatment with the Sema3A inhibitor TCS1105 reversed the delayed fracture healing observed in WT mice following OVX. (A) Schematic of the experimental design to assess the effects of the Sema3A inhibitor TCS1105 on callus remodeling during osteoporotic fractures. (B and G) Representative micro-CT images and quantification of femoral fractures of WT mice treated with TCS1105 or vehicle at 4 weeks post-fracture. Parameters include bone volume fraction (BV/TV). Scale bar: 1 mm. n=6. (C-D) Representative SO-FG (C) and Masson (D) stained images of femoral fractures in WT mice treated with TCS1105 or vehicle at 4 weeks post-fracture. Upper panels display global views; lower panels show close-ups of the fracture sites. Tb: trabecular bone. Scale bar: 1 mm. (E and H) Immunofluorescence staining and quantification of CGRP protein levels in calluses from WT mice treated with TCS1105 or vehicle at 4 weeks post-fracture. ca: callus. White dashed lines outline callus boundaries. Scale bar: 200 μm. n=3. (F, I and J) Immunofluorescence staining and quantification of OCN and CD31 protein levels in calluses from WT mice treated with TCS1105 or vehicle at 4 weeks post-fracture. "ca" denotes callus. White dashed lines outline callus boundaries. Scale bar: 200 μm. n=3. Data presented as mean ± SEM. *P < 0.05, **P < 0.01, ***P < 0.001, ****P < 0.0001.

TCS1105 treatment in OVX mice enhanced bone callus formation and mineralization (Fig. 6 B and Fig.S4 D). Micro-CT analysis revealed that the callus BV/TV in TCS1105-treated mice increased by 61% compared to the vehicle group (Fig. 6 G). Additionally, the number of trabeculae in the bone callus rose by 91%, while trabecular thickness decreased by 15% (Fig.S4 E-G).

We subsequently examined the effects of TCS1105 on new bone formation within the bone callus (Fig. 6 C and D and Fig.S4 H). SO-FG and Masson staining revealed that while new bone formation in OVX mice was reduced compared to sham, it was significantly enhanced in the TCS1105-treated group relative to the vehicle group (Fig. 6 C and D). We also evaluated the impact of TCS1105 on sensory nerve innervation and vascular regeneration. Results indicated a 50% increase in CGRP protein expression in the bone callus post-treatment compared to the vehicle group (Fig. 6 E and H). Similarly, expressions of CD31 and osteocalcin (OCN) were elevated by 45% and 80%, respectively, compared to the vehicle group (Fig. 6 F, I and J). Therefore, targeting Sema3A in the fracture repair microenvironment presents a promising strategy for treating osteoporotic fractures.

### Elevated Cx3cr1^+^iOCs and diminished CGRP^+^TrkA^+^ Signaling in osteoporotic fracture tissue samples

To further assess the expression of Cx3cr1^+^iOCs and CGRP^+^TrkA^+^ signaling in osteoporotic fractures, we conducted histological examinations on a femoral neck specimen from an 88-year-old female undergoing hip replacement surgery (Fig. 7 A). For comparison, a femoral neck specimen from a 55-year-old male patient served as a control (Fig. 7 H). Both patients had surgery three days post-fracture, providing a basis for comparison. HE and SO-FG staining revealed significantly fewer trabeculae in the 88-year-old female compared to the younger male, indicating more severe osteoporosis (Fig. 7 B-C and I-J). More importantly, the expression of Cx3cr1^+^iOCs significantly increased in the subchondral bone of the 88-year-old female, whereas in the younger male, it primarily consisted of normal TRAP^+^ osteoclasts (Fig. S5). Dual immunofluorescence staining for Cx3cr1 and Sema3A showed significantly higher co-localization in osteoporotic fractures compared to the control (Fig. 7 D, K and O). Conversely, the expression levels of CGRP, TrkA, and TUBB3 proteins were markedly lower in the osteoporotic fractures compared to the control group (Fig. 7 F-G, L-N and P-R). These findings suggest a close association between elevated Sema3A expression from Cx3cr1^+^iOCs and diminished CGRP^+^TrkA^+^ signaling at the fracture sites in osteoporotic fractures.

**Figure. 7.**
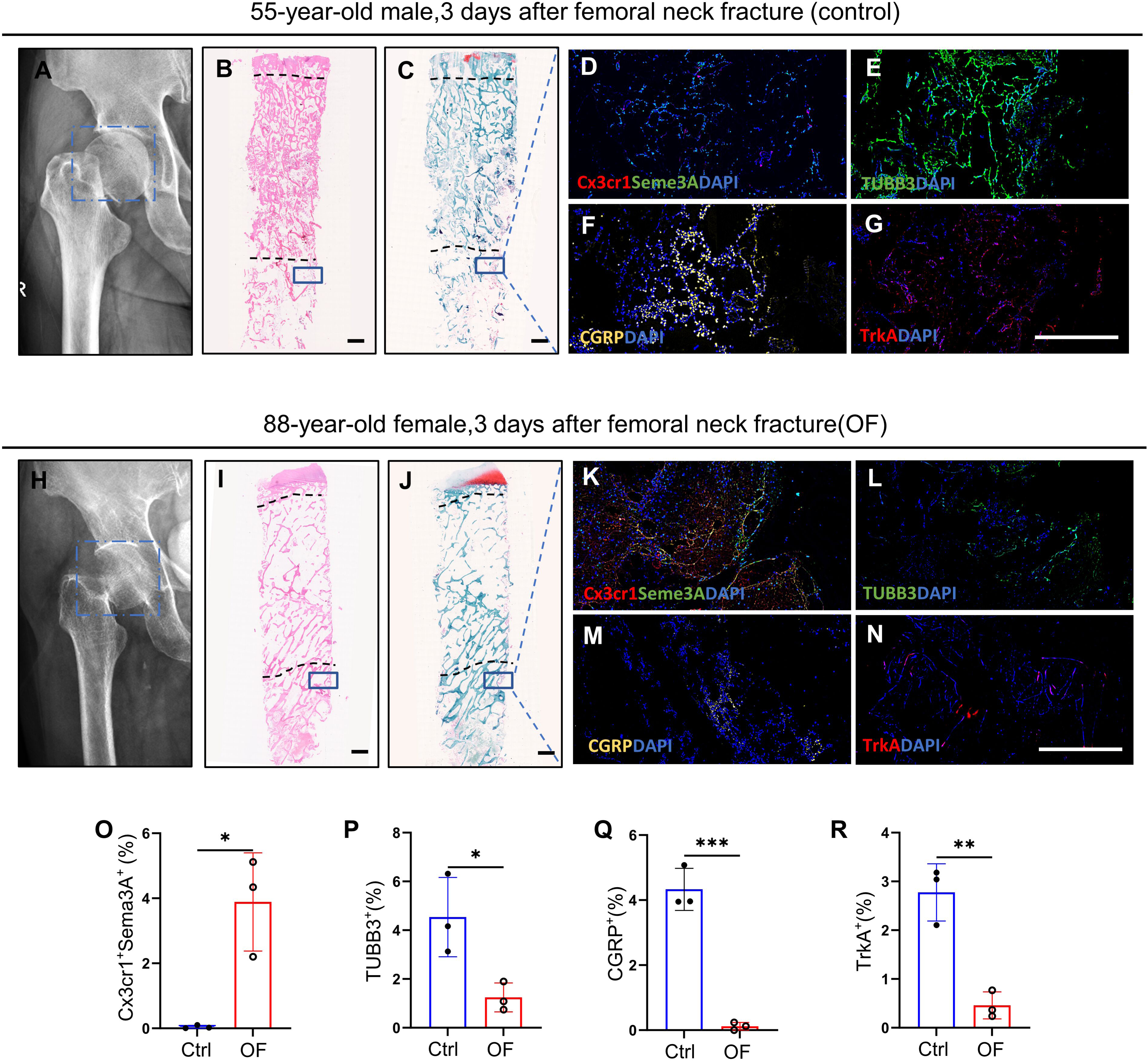
Elevated Cx3cr1^+^iOCs and diminished CGRP^+^TrkA^+^ Signaling in osteoporotic fracture tissues samples. (A-G) Pathological analysis of a femoral neck fracture in a 55-year-old male. Representative X-ray (A), H&E staining (B), and SO-FG staining (C) images are shown on the left. Immunofluorescence images of C x3cr1 and Sema3A coimmunostaining (D), TUBB3 protein (E), CGRP protein (F), and TrkA protein (G) are shown on the right. Scale bar: 1 mm. (H-N) Pathological analysis of a femoral neck fracture in an 88-year-old female. Representative X-ray (H), H&E staining (I), and SOFG staining (J) images are shown on the left. Immunofluorescence images of C x3cr1 and Sema3A coimmunostaining (K), TUBB3 protein (L), CGRP protein (M), and TrkA protein (N) are shown on the right. Scale bar: 1 mm. (O-R) Quantitative analysis of fluorescence intensity for Cx3cr1^+^Sema3A^+^, TUBB3, CGRP, and TrkA in fracture tissue samples from (D-G) and (K-N). n = 3 per group; data presented as mean ± SEM. *P < 0.05, **P < 0.01, ***P < 0.001 and ****P < 0.0001.

## Discussion

Callus remodeling is a critical phase in the fracture repair process. During this phase, osteoclasts not only promote new bone formation through degradation but also participate in angiogenesis and nerve regeneration, completing the final remodeling of the bone(Neto et al., 2022; Liu et al., 2021; Kim et al., 2018; Yang et al., 2012). In the pathological state of osteoporosis, the normal process of callus remodeling is disrupted, significantly contributing to delayed fracture healing. To date, no effective method has been developed to treat the delayed healing of fractures caused by osteoporosis, partly due to a limited understanding of the mechanisms obstructing osteoclast coupling in callus remodeling. In this study, we initially discovered that the loss of CGRP^+^TrkA^+^ sensory neural signaling during callus remodeling in osteoporotic fractures correlates with increased expression of pathological osteoclasts-Cx3cr1^+^iOCs. Subsequently, by specifically deleting Cx3cr1^+^iOCs and Sema3A secreted by Cx3cr1^+^iOCs, we observed enhanced CGRP^+^TrkA^+^ sensory neural signaling within the callus and an acceleration of callus remodeling. Finally, through clinical samples, we confirmed the regulatory relationship between Cx3cr1^+^iOCs and CGRP^+^TrkA^+^ sensory neural signaling (Fig.8).

**Figure. 8.**
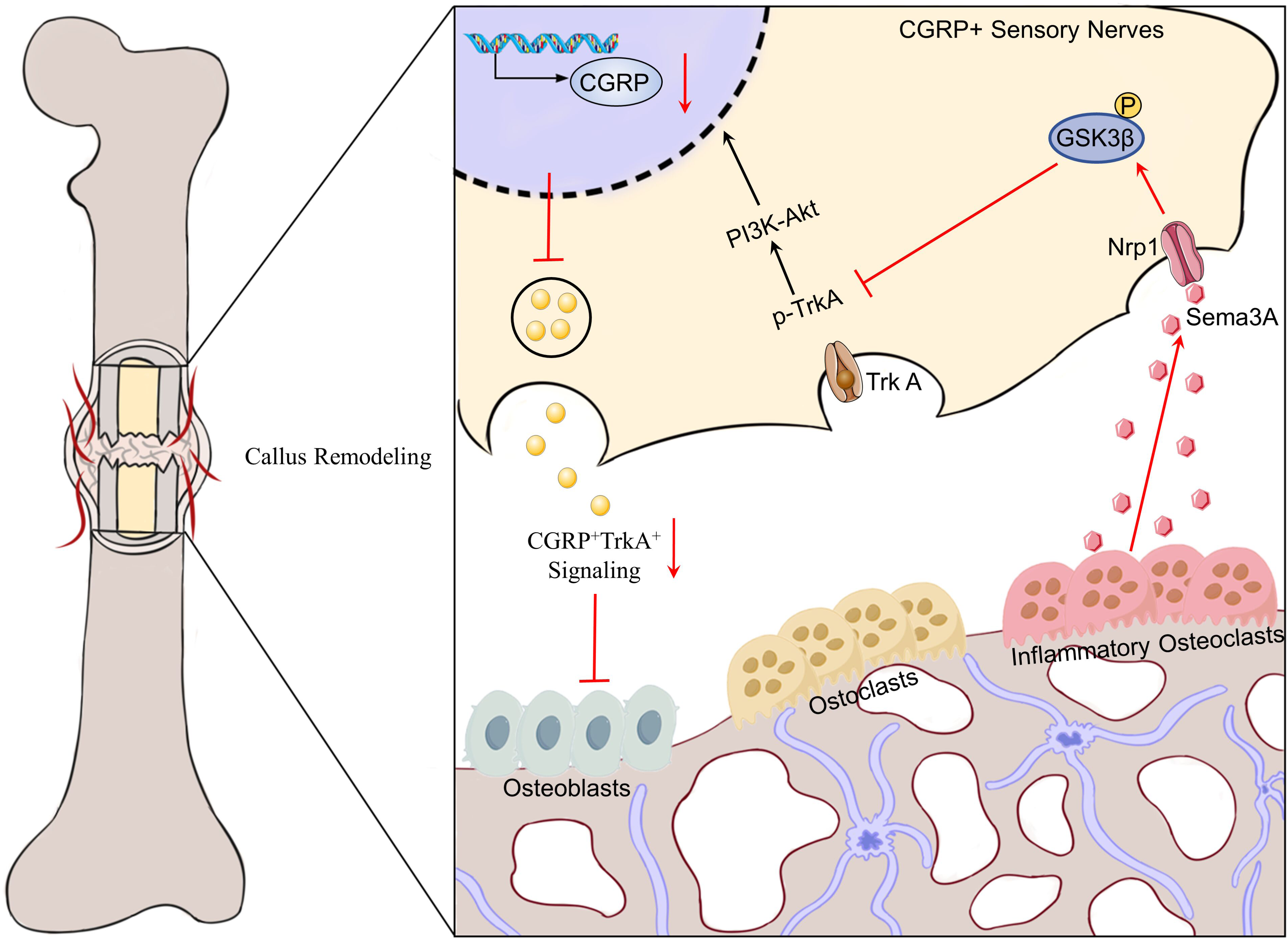
Graphical Abstract: Inhibition of Inflammatory Osteoclasts Enhances CGRP+TrkA+ Signaling and Accelerates Callus Remodeling in Osteoporotic Fractures.

Bone tissue is enveloped by a dense network of sensory nerves, predominantly those expressing calcitonin gene-related peptide (CGRP)(Ni et al., 2019). CGRP is crucial for promoting bone formation, inhibiting bone resorption, inducing angiogenesis, and regulating the immune microenvironment, thereby establishing a vital link between neural activity and bone repair(Wang et al., 2023). Recent studies in a stress fracture model have shown that significantly enhanced CGRP^+^ sensory neural signaling activates tropomyosin receptor kinase A(TrkA) signaling, which is essential for nerve reinnervation, vascular formation, and subsequent osteoblastic activity within the callus(Li et al., 2019). Inhibition of CGRP^+^TrkA^+^ signaling results in delayed callus mineralization(Li et al., 2019). Our study further confirms that in osteoporotic fractures, the absence of CGRP^+^TrkA^+^ signaling significantly delays callus remodeling, underscoring the critical role of sensory nerves in regulating fracture repair. Additionally, we discovered that increased expression of Cx3cr1^+^iOCs within the callus is a key factor in inhibiting sensory neural signaling in the repair microenvironment of osteoporotic fractures.

Callus remodeling is essential for restoring bone integrity and function. Under physiological conditions, osteoclasts support osteoblasts in forming new bone trabeculae by degrading the cartilage matrix, a process essential for callus remodeling(Park et al., 2021). Research indicates that inhibiting osteoclast function may impede bone callus remodeling(Sánchez and Blanco, 2017; Solomon et al., 2009). Osteoclasts, multifunctional cells originating from monocytes and macrophages, exhibit significant heterogeneity across various microenvironments(Guilliams et al., 2018). Our previous studies have demonstrated that osteoclast differentiation, formation, and function are altered under pathological conditions such as inflammation or infection(Ren et al., 2024; Wang et al., 2017). Recent studies have found that in pathological states, osteoclasts known as inflammatory osteoclasts (iOCs), which express high levels of Cx3cr1, show reduced bone resorption and increased immunosuppressive effects(Madel et al., 2020). It remains unclear whether the disruption of callus remodeling in osteoporotic fractures is associated with Cx3cr1^+^iOCs. Our research indicates that the expression of Cx3cr1^+^iOCs in the calluses of osteoporotic fractures significantly increases and that specifically targeting these cells can reverse the disruption of callus remodeling. Given the concurrent loss of CGRP^+^TrkA^+^ signaling, we propose that local neural dysregulation caused by Cx3cr1^+^iOCs may contribute to abnormal callus remodeling.

Next, we explored the molecular mechanisms through which Cx3cr1^+^iOCs contribute to the loss of CGRP^+^TrkA^+^ signaling during callus remodeling. Our findings show that increased expression of Sema3A by Cx3cr1^+^iOCs within the callus inhibited the regeneration of sensory neuronal axons, thereby impeding callus remodeling and resulting in the loss of CGRP^+^TrkA^+^ signaling. Sema3A, an inhibitor of axonal growth, restricts the growth and migration of sensory neuronal axons (Dontchev and Letourneau, 2002). Numerous studies have shown that Sema3A, widely expressed in bone tissue, plays a crucial role in maintaining bone balance and influences osteoblast differentiation through cellular autonomous mechanisms(Fukuda et al., 2013). Recent studies have indicated that under mechanical load, Sema3A secreted by sensory nerves can stimulate bone formation, suggesting that the role of Sema3A in bone repair regulation varies based on its source(Mei et al., 2024). In our study, by specifically deleting Sema3A in Cx3cr1^+^iOCs, we confirmed its role in enhancing sensory neural signaling within the callus, thereby restoring normal neuro-skeletal regulatory mechanisms. Consequently, Sema3A represents a potential therapeutic target for treating delayed healing in osteoporotic fractures. Furthermore, by administering TCS1105 at the fracture site, we were able to enhance CGRP signaling within the callus, promoting both mineralization and angiogenesis.

Ultimately, through the analysis of clinical pathological samples, we discovered that in osteoporotic fractures, the reduction of local CGRP^+^TrkA^+^ signaling is closely associated with increased expression of Cx3cr1^+^iOCs. These findings further substantiate the critical regulatory role of CGRP^+^TrkA^+^ sensory neural signaling in fracture repair and suggest that leveraging CGRP modulation to enhance the interaction between neuro-skeletal systems could be an effective strategy for treating osteoporotic fractures.

Our study has some limitations. Although we demonstrated the interaction between CX3CR1-iOCs and Sema3A impacting CGRP+TrkA+ signaling in callus remodeling through mouse models and observed similar interactions in human tissues, the dynamics of these signals in human fracture repair require further investigation. Additionally, determining the optimal timing and delivery system for administering Sema3A inhibitors in osteoporotic fracture repair remains a challenge. To address this, we are investigating the use of biomaterials such as hydrogels and nanomaterials as carriers to precisely control drug release, thus providing innovative strategies for treatment. Currently, we are developing a hydrogel infused with growth factors that promote callus formation and mineralization, mimicking normal bone repair to facilitate osteoporotic fracture repair. We will continue to explore its effects in future studies.

## Methods and materials

### Mouse models

We acquired genetically modified mice expressing Cx3cr1-Cre, DTR^flox/flox^, and Sema3A^flox/flox^ alleles from Cyagen Biosciences (Suzhou, China), all maintained on a C57BL/6J genetic background. These mice were housed in the Specific Pathogen Free

(SPF) facilities of the Experimental Animal Center at Shanghai Jiao Tong University School of Medicine. The study was approved by the Ethics Committee of Tongren Hospital, affiliated with Shanghai Jiao Tong University School of Medicine (ethical approval number A2023-057-01). All experiments were conducted in accordance with the guidelines of the Experimental Animal Science at Shanghai Jiao Tong University School of Medicine.

To simulate clinical osteoporosis, we established an ovariectomy (OVX) model using the following method: 10-week-old mice were anesthetized with 0.5% pentobarbital sodium administered intraperitoneally, followed by a midline abdominal incision. Abdominal muscles and peritoneum were bluntly separated to locate and ligate the ovaries for removal. The incision was then sutured. The establishment of the osteoporosis model was confirmed two months post-operation.

To investigate fracture healing under osteoporotic conditions, we developed a femoral fracture model in mice post-OVX surgery. Following intraperitoneal anesthesia with 0.5% sodium pentobarbital, the mice’s limbs were secured. Both lower limbs were disinfected with iodine, and a 1cm skin incision was made on the lateral side of the distal femoral condyle. Blunt dissection was used to expose the femur from the intermuscular space fully. A 0.5mm Kirschner wire was retrogradely inserted into the medullary cavity at the femoral condyles, and a stable midshaft fracture was created. Postoperatively, muscles and skin were sutured in layers. The surgical site was disinfected daily, and intramuscular antibiotics were administered until the third day post-operation. At 1-, 2-, 4-, and 8-weeks post-operation, six mice from each group were euthanized for further analysis.

### Human samples

During hip replacement surgeries, we collected human samples from femoral neck fractures. Initially, samples were sectioned from the center of the femoral head to the fracture end into strips approximately 1 cm wide. These samples were then fixed in 4% paraformaldehyde (PFA) solution (G1101-500ML, Servicebio, Wuhan, China) for 48 hours. After fixation, the samples underwent decalcification in 10% EDTA solution (G1105-500ML, Servicebio, Wuhan, China) for six months with continuous agitation. Following decalcification, the samples were embedded in paraffin and sectioned into slices 5 μm thick for further histological analysis. All experimental protocols were approved by the Ethics Committee of Tongren Hospital affiliated with Shanghai Jiao Tong University School of Medicine (ethical approval number 2023-054-01).

### Micro-CT analyses

Femur specimens were imaged using a micro-computed tomography scanner (µCT; Skyscan1276, Bruker microCT, Kontich, Belgium). To prevent any movement during the scanning process, the femurs were securely fastened within the scanning container. Each femur was scanned at settings of 100 kV and 200μA to achieve a voxel resolution of 6 µm. Following scanning, the data were reconstructed using NRecon software. The reconstructed images were then quantitatively analyzed with Bruker’s CTAN software. The volume of interest (VOI) for analysis included the entire healing tissue, while excluding adjacent bone structures. Primary analysis metrics comprised the ratio of healed bone volume to total volume (BV/TV), trabecular thickness (Tb.Th), and trabecular number (Tb.N), structure model index (SMI). All analyses were performed blindly to maintain the objectivity and accuracy of the results.

### Histological staining and immunofluorescence staining

Initially, the mice were anesthetized with 0.5% sodium pentobarbital and secured on the operating table. The femoral blood vessels were then flushed alternately with saline and 4% paraformaldehyde (PFA) solution via cardiac perfusion. Femur samples were subsequently fixed in 4% PFA solution for 48 hours, followed by decalcification in 10% EDTA solution with agitation for four weeks. After decalcification, the samples were embedded in paraffin and sectioned into 5 µm thick slices. These sections were stained with hematoxylin and eosin (H&E), Masson’s trichrome, and saffron O/fast green (SOFG) to analyze the remodeling within the callus.

For immunofluorescence staining, the paraffin sections were first dewaxed twice in xylene, each time for 10 minutes. Sections were then dehydrated through a graded ethanol series (100%, 95%, 80%, and 75%), each concentration for 5 minutes. Antigen retrieval was performed in citrate buffer (pH 6.0) using a microwave: heating initially at high power for 8 minutes, pausing for 8 minutes, and then continuing at medium-low power for 7 minutes. After cooling, sections were blocked with 3% bovine serum albumin (BSA) for 30 minutes. The primary antibody was diluted to the recommended concentration and incubated at 4°C overnight. Following primary antibody incubation, sections were washed three times with PBS, then incubated with a diluted secondary antibody (1:1000) at room temperature for 60 minutes. Nuclei were stained with 4’,6-diamidino-2-phenylindole (DAPI) and visualized under an Olympus microscope. Positive cells or areas in the sections were quantitatively analyzed using ImageJ software.

### Co-culture under cell conditions

#### Primary isolation of BMMs

Initially, monocytes and macrophages (BMMs) are harvested from the bone marrow cavities of mouse femurs and tibias. These cells are then cultured overnight in αMEM (Gibco, USA) medium enriched with 10% fetal bovine serum (Gibco, USA), 1% penicillin/streptomycin, and 30 ng/ml macrophage colony-stimulating factor (M-CSF) (R&D Systems, MN). After the removal of adherent cells, the non-adherent cells are cultured further in medium containing 30 ng/ml M-CSF to isolate pure BMMs. Finally, the BMMs are incubated for three days in a 24-well plate with medium supplemented with 30 ng/ml M-CSF and 50 ng/ml receptor activator of nuclear factor kappa-B ligand (RANKL) (R&D Systems, MN). After this period, nearly all the cells have differentiated into osteoclasts.

#### Sema3A shRNA construction

Sema3A shRNA was sourced from Cyagen Biosciences (Suzhou, China). The sequences of shRNA-Sema3A employed were sense strand, 5′-AATTCAAAAATGCCAGTTTGTCAAATACCCCA-3′; and antisense strand, 5′-CCGGTGGTTCATTTGACAAACTGACATGCATTTG-3′. Cells were transfected using Lipofectamine 2000 (Life Technologies, USA) following the manufacturer’s instructions. After 72 hours of incubation, cell supernatants were collected, centrifuged at 2500 rpm for 10 minutes at 4°C, and the conditioned mediums were aliquoted and stored at −80°C. Isolation and culture of DRG neurons The isolation and culture of dorsal root ganglia (DRGs) neurons are conducted according to the protocol by Patrick et al.(Smith et al., 2023). Specific steps include selecting 4-week-old mice, which are then euthanized following anesthesia. In a sterile environment, the spine is carefully dissected under a stereomicroscope to extract the dorsal root ganglia. Immediately after extraction, the ganglia are placed in precooled Hank’s Balanced Salt Solution (HBSS) (Gibco, USA) containing 2% penicillin/streptomycin. Subsequently, the DRGs are incubated at 37°C with 1 mg/mL collagenase A (10103578001, Roche) for 15 minutes, followed by a 10-minute digestion with 0.05% trypsin-EDTA at the same temperature. Post-digestion, cells are separated by centrifugation and resuspended in complete DRG medium. The cells are then seeded onto culture plates coated with poly-D-lysine (PDL) (Coolaber, China). Six hours post-adherence, the medium is replaced with fresh complete Neurobasal medium (Gibco, USA). Cells are monitored and imaged using a fluorescence microscope for subsequent experiments.

### Flow Cytometry

First, bone marrow is extracted from the femoral marrow cavity of a mouse, and a 70-μm nylon mesh is used to filter out debris. The filtered cell suspension is then transferred into a 1.5 mL centrifuge tube and centrifuged at 1500 rpm for 5 minutes to pellet the cells. After centrifugation, the cells are resuspended in 1 mL of precooled PBS and centrifuged again at 1500 rpm for 5 minutes; the supernatant is subsequently discarded. Following this, the cells are stained with specific fluorescent dyes using antibodies targeting Cx3cr1 (1:100, NBP2-68616, Novus) and Ly6C (1:100, 88920, Cell Signaling Technology). Once stained, the cells are analyzed on a flow cytometer using the appropriate channels. FlowJo software is employed to analyze the data, focusing on calculating the positivity rate and average fluorescence intensity of the positive regions to elucidate the phenotype and functional state of the cells.

### Western blot

Add an appropriate volume of RIPA lysis buffer (G2002-30ML, Servicebio, China) to the DRGs cell culture plate and incubate on ice for 30 minutes to lyse cells. Centrifuge at 12,000 rpm and 4°C for 15 minutes, then collect the supernatant to obtain the total protein solution. After measuring the protein concentration, add 5X reducing sample buffer and heat at 95°C for 10 minutes to denature the proteins. Perform SDS-PAGE for protein electrophoresis. Transfer the proteins onto a PVDF membrane (ISEQ00010, Merck Millipore, Germany). Block the membrane with 5% skim milk for 30 minutes, followed by incubation with the primary antibody at 4°C for 12 hours. Wash the membrane three times with TBST, then incubate with a 1:1,000 dilution of the secondary antibody (7074, Cell Signaling Technology) at room temperature for one hour. After further washes, detect chemiluminescent signals with the Bio-Rad Molecular Imager ChemiDocTM XRS+ system (Bio-Rad, USA). Quantitatively analyze the protein expression using ImageJ software. The following antibodies were used: CGRP (1:500, 14959S, Cell Signaling Technology), TrkA (1:500, 2510, Cell Signaling Technology), p-TrkA (1:500, 9141, Cell Signaling Technology), Akt and p-Akt (1:1000, 4060T, Cell Signaling Technology), GSK3β and p-GSK3β (1:500, AF1590-SP, R&D Systems), GAPDH (1:500, 8884, Cell Signaling Technology).

### Statistical analysis

Data are presented as mean ± SD or SEM. Two-group comparisons were conducted using the two-tailed Student’s t-test. Multiple group comparisons were analyzed via one-way ANOVA. Results were visualized and analyzed using GraphPad PRISM 9.0, with P < 0.05 indicating statistical significance.

## Acknowledgments

This work was supported by a grant from the National Natural Science Foundation of China (82102577), the Laboratory Open Fund of Key Technology and Materials in Minimally Invasive Spine Surgery (2024JZWC-YBA05;2024JZWC-YBA01), the Tongren Hospital Introduces the Talented Person Scientific Research Start Funds Subsidization Project (TR2023rc08).

## Author contributions

Yuan Wang designed the study and drafted the manuscript. Yuexia Shu and Zhenyu Tan conducted the majority of experiments. Zhen Pan, Yujie Chen, and Jielin Wang performed statistical analyses. Jieming He and Jia Wang collected clinical specimens. All authors approved the final manuscript.

## Data Availability Statement

Data related to this study are presented within the main text, figures, and supplementary materials. All data can be obtained from the corresponding author upon reasonable request.

**Figure.S1. Reduced bone mass and delayed fracture healing in OVX mice.**

(A) Representative Micro-CT images of femurs in sham and OVX mice. Scale bar: 1 mm.

(B-D) Micro-CT quantitative analysis of the femurs from (A). The Micro-CT parameters include bone volume fraction (BV/TV) of calluses, Tb.Th (trabecular thickness), Tb.N (trabecular number). n=3.

(E) Representative micro-CT images of femoral fractures at 2- and 4-weeks post-fracture.

(F) Micro-CT quantitative analysis of the callus from (E). The micro-CT parameters include the structure model index (SMI). n=6.

(G) Representative images of Masson staining of femoral fractures in sham and OVX mice at 2- and 4-weeks post-fracture. Upper panels display global views; lower panels show close-up views of the fracture sites. Tb: trabecular bone. Scale bar: 1 mm. Data presented as mean ± SEM. *P < 0.05, **P < 0.01, ***P < 0.001, ****P < 0.0001.

**Figure.S2. Conditional knockout of CX3CR1^+^iOCs accelerates callus remodeling by enhancing CGRP+TrkA+ signaling.**

(A) Representative micro-CT images of femoral fractures of Cx3cr1-Cre; DTR^fl/fl^ mice and WT littermates at 4 weeks post-fracture.

(B) Representative images of Masson staining of the femoral fractures of Cx3cr1-Cre; DTR^fl/fl^ mice and WT at 4 weeks post-fracture. The upper panels display global views, while the lower panels show close-up views of the fracture sites. Tb: trabecular bone. Scale bar: 1 mm.

(C) Immunofluorescence staining images showing TUBB3 protein levels in calluses from Cx3cr1-Cre; DTR^fl/fl^ mice mice and WT littermates at 2 weeks post-fracture. ca: callus. Scale bar: 200 μm. Data presented as mean ± SEM. *P < 0.05, **P < 0.01, ***P < 0.001, ****P < 0.0001.

**Figure.S3. Sema3A secreted by Cx3cr1^+^iOCs inhibits CGRP^+^TrkA^+^ sensory nerve signaling via GSK3β-Akt pathway in vitro.**

(A) Flow cytometry analysis of Cx3cr1^+^Ly6C^+^ cells in sham and OVX mouse bone marrow. n=3.

(B) Representative coimmunostaining images of CX3CR1, Sema3A, Sema4D, and Sema7A, and quantification in calluses from sham and OVX mice at 4 weeks post-fracture. ca: callus. white arrows point to Cx3cr1^+^ Sema3A^+^. Scale bar: 200 μm.

(C and E) Western blot analysis of Akt and p-Akt in DRGs treated with or without Sema3A from by cx3cr1^+^iOCs.

(D) Western blot analysis of CGRP, TrkA, p-TrkA in DRGs treated with Sham, OVX or OVX+anti-GSK3β.

(F-G) Quantitative analysis of CGRP, TrkA and p-TrkA expression from (D). n = 3 per group; data presented as mean ± SEM. *P < 0.05, **P < 0.01, ***P < 0.001 and ****P < 0.0001.

**Fig.S4. Targeting Sema3A in Cx3cr1^+^iOCs enhances CGRP^+^TrkA^+^ signaling and promotes callus remodeling.**

(A) Representative SO-FG-stained images of femoral fractures in Cx3cr1-Cre; Sema3A^fl/fl^ OVX mice and their littermate controls at 4 weeks post-fracture. Upper panels display global views; lower panels show close-up views of the fracture sites. Tb: trabecular bone. Scale bar: 1 mm.

(B-C) Immunofluorescence staining images and quantification of TUBB3 protein levels in calluses from Cx3cr1-Cre; Sema3A^fl/fl^ OVX mice and their littermate controls at 4 weeks post-fracture. ca: callus. White dashed lines outline the callus boundaries. Scale bar: 200 μm. n=3.

(D) Representative micro-CT images of femoral fractures of WT mice treated with TCS1105 or vehicle at 4 weeks post-fracture. Scale bar: 1 mm.

(E-G) Micro-CT quantitative analysis of the callus from (D). Parameters include trabecular thickness (Tb.Th), trabecular number (Tb.N), and structure model index (SMI). n=6.

(H) Representative images of H&E staining of femoral fractures in WT mice treated with Sema3A inhibitor or normal saline at 4 weeks post-fracture. Upper panels display global views; lower panels show close-up views of the fracture sites. Tb: trabecular bone. Scale bar: 1 mm. Data presented as mean ± SEM. *P < 0.05, **P < 0.01, ***P < 0.001, ****P < 0.0001.

**Fig.S5. Elevated Cx3cr1^+^iOCs and diminished CGRP^+^TrkA^+^ Signaling in osteoporotic fracture tissue samples.**

(A-D) Pathological analysis of a femoral neck fracture in a 55-year-old male. Representative images of Masson staining (A) and detailed images (B) on the left. Immunofluorescence staining of CX3CR1 and TRAP coimmunostaining (C), and detailed images (D) on the right. Scale bar: 1 mm.

(E-H) Pathological analysis of a femoral neck fracture in an 88-year-old female. Representative images of Masson staining (E) and detailed images (F) on the left. Immunofluorescence staining of CX3CR1 and TRAP coimmunostaining (G), and detailed images (H) on the right. Scale bar: 1 mm.

(I) Quantitative analysis of CX3CR1 and TRAP fluorescence intensity in fracture tissue samples from (C) and (G). n = 3 per group; data presented as mean ± SEM. *P < 0.05, **P < 0.01, ***P < 0.001 and ****P < 0.0001.

**Figure.S6. Western blot original bands and ethical approval documentations**

(A) Western blot original bands.

(B) ethical approval documentations

